# B7-H3-targeted natural killer cells effectively kill atypical teratoid / rhabdoid tumors and extend survival in orthotopic xenografts

**DOI:** 10.64898/2026.01.15.699746

**Authors:** Jun Choe, Sachiv Chakravarti, Natalie J. Holl, Megan T. Zinsky, Danielle G. Jones, Shramana Guchhait, Ruyan Rahnama, Stamatia C. Vorri, Adrianna Amaral, Calixto-Hope G. Lucas, Eric H. Raabe, Challice L. Bonifant

**Author notes:** Corresponding Author: Challice L. Bonifant, MD, PhD,; Eric H. Raabe, MD, PhD, 1550 Orleans St., Baltimore, MD 21287. Declaration of interests C.L.B. has been awarded and has pending patent applications describing the use of engineered T and NK cells as therapeutics. C.L.B. has received research support from Merck, Sharp, and Dohme, Inc, and Bristol-Myers Squibb.

## Abstract

**Background:** Atypical teratoid/rhabdoid tumors (AT/RTs) are the most common malignant CNS tumor in infants, and patients suffer from low survival rates and treatment-related morbidities. These tumors frequently overexpress the pan-cancer antigen B7-H3 (CD276), which can be targeted with immunotherapy. We hypothesized that adding a B7-H3-targeting cytotoxic chimeric antigen receptor (CAR) to NK cells can enhance killing against AT/RTs.

**Methods:** We designed a library of variable affinity B7-H3-targeted CARs, which were transduced into primary healthy donor-derived NK cells. We verified B7-H3 expression in a panel of AT/RT cell lines and further engineered luciferase and nuclear GFP-expressing AT/RT (CHLA-04, CHLA-06, BT12, BT37) as well as a CHLA-06-derived B7-H3 knockout. We tested CAR-NK cell functionality using *in vitro* co-culture cytotoxicity assays. We delivered anti-B7-H3 CAR-NK cells intratumorally or intracerebroventricularly (ICV) to AT/RT orthotopic xenografts and monitored for tumor growth and animal survival.

**Results:** B7-H3-targeted CAR-NK cells demonstrated target-specific cytotoxicity when compared to unmodified NK cells. Knockout of B7-H3 in target cells abolished the increased CAR-mediated target killing. When delivered intratumorally to CHLA-06 orthotopic xenograft-bearing mice, anti-B7-H3 CAR-NK cells eliminated tumor cells and prolonged survival. When CAR-NK cells were delivered ICV against a CNS disseminated tumor model of BT12, treated mice had significantly improved survival.

**Conclusions:** Anti-B7-H3 CAR-NK cells effectively kill AT/RTs in multiple pre-clinical *in vitro* and *in vivo* models in an antigen-specific manner. Evidence of efficacy in translationally relevant models provides support for using B7-H3-targeting CAR-NK cells in high-risk AT/RT patients.

## Introduction

Atypical teratoid / rhabdoid tumors (AT/RTs) are aggressive pediatric central nervous system (CNS) cancers that arise most commonly in infants^1^. AT/RTs are characterized by loss-of-function mutations in *SMARCB1*, and are separated into three molecular subtypes (SHH, MYC, TYR)^2^. Despite aggressive multimodal therapies, patients suffer from high relapse rates and treatment related morbidities, emphasizing a need for novel targeted therapies^3^. AT/RTs and other pediatric brain tumors have high expression of the pan-cancer antigen B7-H3 (encoded by *CD276*)^4,5^. Studies have shown effective targeting of B7-H3 using immunotherapy including antibodies^6^, antibody-drug conjugates (ADC)^7^, radiolabeled antibodies^8^, and chimeric antigen receptor (CAR) T cells^9–12^.

B7-H3 is a member of the B7 family of immunomodulatory proteins and has both stimulatory^4^ and inhibitory^13–17^ effects on lymphocytes, although inhibition is more prominent. B7-H3 expression positively correlates with tumor aggressiveness^18^, but its causative role and mechanism of action have not been fully elucidated. Structurally, human B7-H3 protein is composed of two duplicated sets of the extracellular immunoglobulin-like (Ig) domains IgV and IgC (4Ig)^19,20^. The mouse counterpart retains a single set (2Ig)^21^. Although human tissue and tumors express 2Ig CD276 mRNA, evaluation of protein expression suggests that 4Ig is the predominantly expressed isoform^22^.

CAR-T cells are currently under clinical investigation in many adult and pediatric brain tumor types^11,23,24^. Pre-clinical validation of B7-H3 and Claudin 6 as CAR-T cell targets for AT/RT has also been shown^10,25^. However, there are limitations to T-cell therapies, including the requirement of patient autologous T cells to avoid the risk of graft-versus-host disease (GVHD). CAR-T cells are also associated with a high prevalence of inflammatory toxicities such as cytokine release syndrome (CRS), immune-effector cell associated neurotoxicity (ICANS), and tumor inflammation-associated neurotoxicity (TIAN)^26–28^. Natural Killer (NK) cells are an “off-the-shelf” alternative to T cells for use in cancer therapy due to a potential for safe allogeneic transfer from healthy donors^29^. NK cells express a variety of germline-encoded cytotoxicity-mediating receptors that respond to activating ligands expressed by cancer cells^30^. This is an advantage when combating tumors that have the potential to downregulate their CAR target and/or HLA receptors. It is also beneficial when tumors exhibit intratumoral heterogeneity. In clinical trials, NK cells have had favorable toxicity profiles, with limited inflammatory adverse events noted secondary to NK-cell infusions^31–33^.

Treatment of CNS tumors with engineered cell therapies has proven challenging, and various systemic and local administration routes have been tested^10,23^. Local delivery either into the cerebrospinal fluid (CSF) or directly into the tumor is the most direct way to bypass the blood-brain-barrier (BBB), and clinical trials have established the safety of T- and NK-cell therapies delivered this way in pediatric patients^11,32^. Here, we generated B7-H3-targeted CARs with a range of single chain variable fragment (scFv) affinities. All constructs expressed efficiently in primary healthy donor-derived NK cells and demonstrated preclinical efficacy against relevant models of AT/RT.

## Materials and methods

Methods for cell culture and reporter cell generation are in supplementary methods.

### CD276 knockout cell line generation

CD276/B7-H3 expression was eliminated from parental CHLA-06 cells by electroporation of a ribonucleoprotein (RNP) consisting of Cas9 enzyme loaded with three single guide RNAs (sgRNAs) according to the manufacturer’s protocol (spCas9 2NLS Nuclease, Synthego). sgRNA sequences were designed to cause an early truncation in exon 2 of human *CD276* (Supplementary Table 1). In short, an RNP complex was formed by mixing and incubating a 1:9 molar ratio of Cas9:sgRNAs at room temperature for 10 minutes. Cells were prepared at 2e5 cells/20µL/well in a 16-well cuvette according to manufacturer’s instructions (SF Cell Line 4D-Nucleofector X Kit S, Lonza). 7 µL of RNP solution was mixed into the cell suspension and electroporated using the DS-138 protocol (4D-Nucleofector X Unit, Lonza). Cells were then transferred to a 12-well tissue culture plate and incubated in complete neurobasal medium. Loss of B7-H3 expression was evaluated by flow cytometry, and B7-H3(-) single cell clones were isolated (BD FACSMelody). Surface B7-H3 negativity was verified by flow cytometry, and genomic DNA was isolated (DNEasy Blood & Tissue, Qiagen) for *CD276* mutation verification by Sanger sequencing (Johns Hopkins Genetic Resources Core Facility). Once knockout clones were validated, all viable clones were pooled into a stable stock at equal ratios.

### Anti-B7-H3 CAR transgene generation

CAR transgenes were designed using a previously described humanized and affinity matured B7-H3-targeting scFv (Hu8H9)^34^. Transgenes were synthesized (GeneArt, ThermoFisher) and subcloned using In-Fusion cloning (Takara Bio) into a previously described pSFG plasmid backbone containing 2B4 hinge (H), transmembrane (TM), and intracellular (IC) domains followed by a CD3ζ IC domain^35^. scFv variants were generated by site directed mutagenesis (SDM) on the parental Hu8H9 scFv (Agilent). A detailed list of primers used for cloning and SDM are in Supplementary Table 1. Subcloned plasmids were amplified in Stellar chemically competent *Escherichia coli* (Takara) and isolated using DNA Mini or Midi kits (Qiagen) following the manufacturer’s instructions. Sequence fidelity was verified by Sanger sequencing (Johns Hopkins Genetic Resources Core Facility). A detailed list of plasmids generated in this study is in Supplementary Table 2.

### Viral vector production

Hu8H9 CAR vectors were produced by plating 3e6 293Vec-BaEV producer cells^36^ in a 10 cm tissue culture dish in Iscove’s Modified Dulbecco’s Medium (IMDM, Gibco) supplemented with 10% FBS and allowed to adhere overnight. Cells were then transfected with GeneJuice Transfection Reagent (Sigma-Aldrich) and 10 µg of total CAR plasmid DNA. Supernatant was collected after 48 hours, passed through 0.45 µm syringe filters, and snap frozen before storing at -80°C. Vectors for a NLS-GFP reporter were produced similarly using HEK293T cells, except the cells were transfected with 10 µg of total plasmid DNA at a ratio of 3:3:2 of packaging genes:VSV-G envelope:NLS-eGFP.

### CAR-NK cell production

Peripheral blood mononuclear cells (PBMCs) were isolated from fresh donor blood by layering over Lymphoprep (STEMCELL Technologies) and centrifuging. Buffy coat was collected, washed, and depleted of T cells using human CD3 MicroBeads (Miltenyi Biotec 130-050-101). CD3 depletion was verified by flow cytometry using fluorophore-conjugated antibodies against CD56 and CD3. A detailed list of antibodies used is in Supplementary Table 3. CD3-depleted PBMCs were stimulated on Day 0 by co-culturing at a 1:1 ratio with lethally irradiated K562 feeder cells^37^ expressing membrane bound IL-21 and 4-1BBL.. Cells were maintained in Advanced RPMI medium (aRPMI) supplemented with 10% FBS, 2 mmol/L GlutaMax (Gibco 35050061), 200 IU/mL human IL-2 (hIL-2, Biological Resources Branch Preclinical Biorepository, National Cancer Institute), and 2 ng/mL human TGF-β1 (R&D Systems 7754-BH). NK cells were transduced on Day 4 of culture using transiently produced replication-incompetent BaEV-pseudotyped retroviral particles immobilized on RetroNectin (Takara T100B). Transduced NK cells were replated onto 24-well tissue culture-treated plates on Day 6, and surface CAR expression was verified between Days 9-12. NK cells were restimulated with irradiated feeder cells at a 1:1 ratio every 7 days (Day 7, 14, 21, etc.). Functional assays were carried out post-verification of CAR positivity and at least 3 days after restimulation with feeder cells. NK cells for all *in vivo* studies were cryopreserved in 90% growth media + 10% DMSO 13 days post activation and thawed in warmed growth media + 50 ug/mL DNase I before resuspension in PBS and subsequent injection into mice.

### Flow cytometry

All samples were acquired on a FACSCelesta Cell Analyzer (BD Biosciences) and analyzed with FlowJo software (v10.10.0). All cell sorting was performed using a FACSMelody (BD Biosciences). All antibody stains were performed in phosphate buffered saline (PBS) + 1% FBS and incubated on ice for 30 minutes in the dark. Live-dead stains were performed in PBS, incubated on ice for 30 minutes in the dark before antibody staining. B7-H3 surface staining on AT/RT cell lines was performed using anti-human B7-H3-APC (BioLegend, MIH42 clone). CAR expression analysis was performed by primary staining of NK cells with His-tagged recombinant 4Ig B7-H3 protein (Sino Biological 11188-H08H) followed by secondary staining with anti-His-PE or anti-His-APC (BioLegend). Titration of recombinant B7-H3 was performed to ensure saturating conditions. Human Fc receptors were blocked using Human Fc block (BD Biosciences 564219). B7-H3 antigens per cell on AT/RT cell lines were quantified using Molecules of Equivalent Soluble Fluorochrome (MESF) microspheres (Bangs Laboratories, 823) according to the manufacturer’s instructions.

### NK Degranulation assay

NK cells were collected after at least 6 days post restimulation with feeder cells and cytokine starved in complete aRPMI overnight. NK cells and CHLA-06 target cells were plated in flat 96-well tissue culture plates at 1:1 effector-to-target (E:T) ratio and incubated at 37°C for 2 hours (200k:200k cells/well/200uL growth media). Cells were collected on ice for live/dead staining followed by surface staining with anti-CD45 and anti-CD107a antibodies then acquired by flow cytometry.

### SDS-PAGE and Western blot (WB)

Cell lysates were collected in ice cold RIPA buffer (Millipore Sigma, R0278) supplemented with 1X protease/phosphatase inhibitor cocktail (ThermoFisher, 78440) and quantified usin a Pierce BCA kit (ThermoFisher 23225). For deglycosylated lysate, 40µg of whole cell lysate was digested using PNGase F following the manufacturer’s instructions (New England Biolabs, P0704). Samples were loaded and run in 10% Bis-Tris polyacrylamide gels (ThermoFisher, NW00100) in 1X MOPS buffer (ThermoFisher, B0001) and transferred onto PVDF membranes using 1X transfer buffer (ThermoFisher BT0006) + 20% methanol. Membranes were blocked with 5% BSA in Tris buffered saline + 0.1% tween 20 (TBST), 5% bovine serum albumin (BSA), and incubated with primary antibodies at 4°C overnight. Secondary HRP-conjugated antibodies and primary anti-GAPDH incubations were performed for 1 hour at room temperature. All membranes were imaged using a BioRad ChemiDoc imager.

### Histology and Immunohistochemistry (IHC)

All whole brain and spine tissue were fixed in 10% neutral buffered formalin for 48-72 hours and stored in 70% ethanol at 4°C until processing. All histology and H&E staining from animal experiment tissue was performed by Johns Hopkins Oncology Tissue & Imaging Services. Spine tissue was decalcified in 10% EDTA, pH 7.4 at 4°C with rocking for 6 days and cross-sectioned axially into ∼2-3mm slices before paraffin embedding and sectioning at 4µm.

Formalin-fixed paraffin-embedded (FFPE) primary tumor tissue was obtained from Johns Hopkins Biobank and stained for human B7-H3 as previously described^38^. Briefly, 4 µm sections were stained with an anti-human B7-H3 primary antibody (clone D9M2L, Cell Signaling Technology 14058) on the Ventana Benchmark immunostaining system (Roche Diagnostics). CC1 was used for antigen retrieval before primary antibody incubation for 1 hour. OptiView DAB (Roche 760-700) was used for detection and hematoxylin and bluing reagent for counterstaining. Intensity and extent of B7-H3 immunoreactivity were scored by a board-certified neuropathologist on a semi-quantitiave basis. Intensity was scored as 1=minimal/weak, 2=moderate, 3=strong, whereas extent was scored as 1=absent/focal, 2=patchy (less than ∼50% examined area), or 3=extensive (greater than ∼50% examined area). The total score was calculated as a product of intensity and extent scores. For evaluation of tumor area and human CD45 quantification from murine samples, areas of viable tumor were annotated. In each case containing viable tumor, the highest absolute count of intra/peritumoral human CD45-positive cells in a single 40x microscopic field was quantified. In each case, the cross-sectional area of the largest tumor focus on a 4x microscopic field was quantified in pixel2 using the polygon tool in QuPath software (v0.3.2)^39^.

### Short-term in vitro cytotoxicity assay

NK cells were cytokine-starved overnight in complete aRPMI. Firefly luciferase (ffLuc)-expressing AT/RT cell lines (CHLA-06, CHLA-06 B7-H3KO, BT12, BT37, CHLA-04) were plated in co-culture with NK cells at indicated E:T ratios (1e5 target cells/well). After 18 hours of incubation, D-luciferin (ThermoFisher Scientific) was added at a final concentration of 150 µg/mL and radiance was measured using a BMG CLARIOstar microplate reader.

### Incucyte live cell imaging serial killing assay

NK cells were cytokine-starved overnight in complete aRPMI. NLS-eGFP expressing target AT/RT cell lines were plated in co-culture with NK cells at a 1:1 E:T ratio (5e4 target cells) in 200 µL/well of complete aRPMI supplemented with a final concentration of 20 IU/mL of hIL-2 onto flat-bottom, poly-D-lysine-coated 96-well plates in triplicate. All co-culture wells were replenished with 5e4 target cells and 20 IU/mL hIL-2 every 48 hours. Phase contrast and green-fluorescent images were captured every 4 hours, with acquisition of 4 images per well.

### In vivo orthotopic xenograft models and treatments

Animal experiments were done in accordance with protocol approved by Johns Hopkins Animal Care and Use Committee (IACUC). All mice were obtained from The Jackson Laboratory or from an internal colony that originated from The Jackson Laboratory. Orthotopic xenografts of human AT/RT cell lines were established in NOD.Cg-*Prkdc^scid^ Il2rg^tm1Wjl^*/SzJ (NSG) mice as previously described^40^. In short, a burr hole was made using an 18-gauge needle (from bregma: 3 mm posterior, 2 mm anterior, 3.5 mm deep for caudate putamen / 0.5 mm posterior, 1 mm anterior, 2.8 mm deep for ICV). A guide-screw (Protech 8L000121207F) was installed into the hole, flush with the skull. A 10 µL 26-gauge Hamilton syringe fitted with a stopper for appropriate depth was used for all injections. 2.5e4 CHLA-06.ffLuc cells/3 µL of Neurobasal medium into the caudate putamen or 1.0e5 BT12.ffLuc cells/3 µL of serum-free RPMI ICV was injected slowly into the bore of the guide-screw before closing with a dummy cannula. Intratumoral and ICV NK cell treatments were performed similarly by accessing the guide-screw with a small incision and removing the dummy cannula. 3.0e6 CAR-NK or unmodified NK cells/5 µL of PBS were injected slowly into the bore of the guide-screw using the same coordinates.

### Statistical analysis

All statistical analyses were performed using GraphPad Prism (v10.4.2). For comparisons between two groups, two-tailed t-tests were used. For comparisons including more than two groups, simple two-way analysis of variance was performed by Dunnett’s multiple comparison test with UTD NK cells used as control group. P-values are displayed as * (≤0.05), ** (≤0.01), *** (≤0.001), and **** (≤0.0001). For long-term cytotoxicity data, area under curve (AUC) values were calculated from normalized live target values at each 48-hour stimulation timepoint. Each AUC value was plotted with its 95% confidence interval shown.

## Results

### All AT/RT molecular subgroups express surface B7-H3

FFPE AT/RT patient tissue acquired from the Johns Hopkins Biobank was evaluated for B7-H3 expression by IHC, with overall moderate-to-high amounts of positivity observed (Figure 1A,B, Supplementary Table 4). For *in vitro* analysis, we chose representative patient-derived cell lines from each of the three AT/RT molecular subgroups (SHH - ATRT310, CHLA-04, CHLA-05; TYR - BT37; MYC - BT12, CHLA-06)^2^. We analyzed surface expression of B7-H3 on select cell lines by flow cytometry. All cell lines stained positively for surface B7-H3, compared to isotype controls and an isogenic CHLA-06 *CD276* knockout (Figure 1C, Figure S1A). Furthermore, antigen density was quantified; all tested cell lines express ∼5,000-20,000 B7-H3 antigens per cell (Figure 1D).

**Figure 1.**
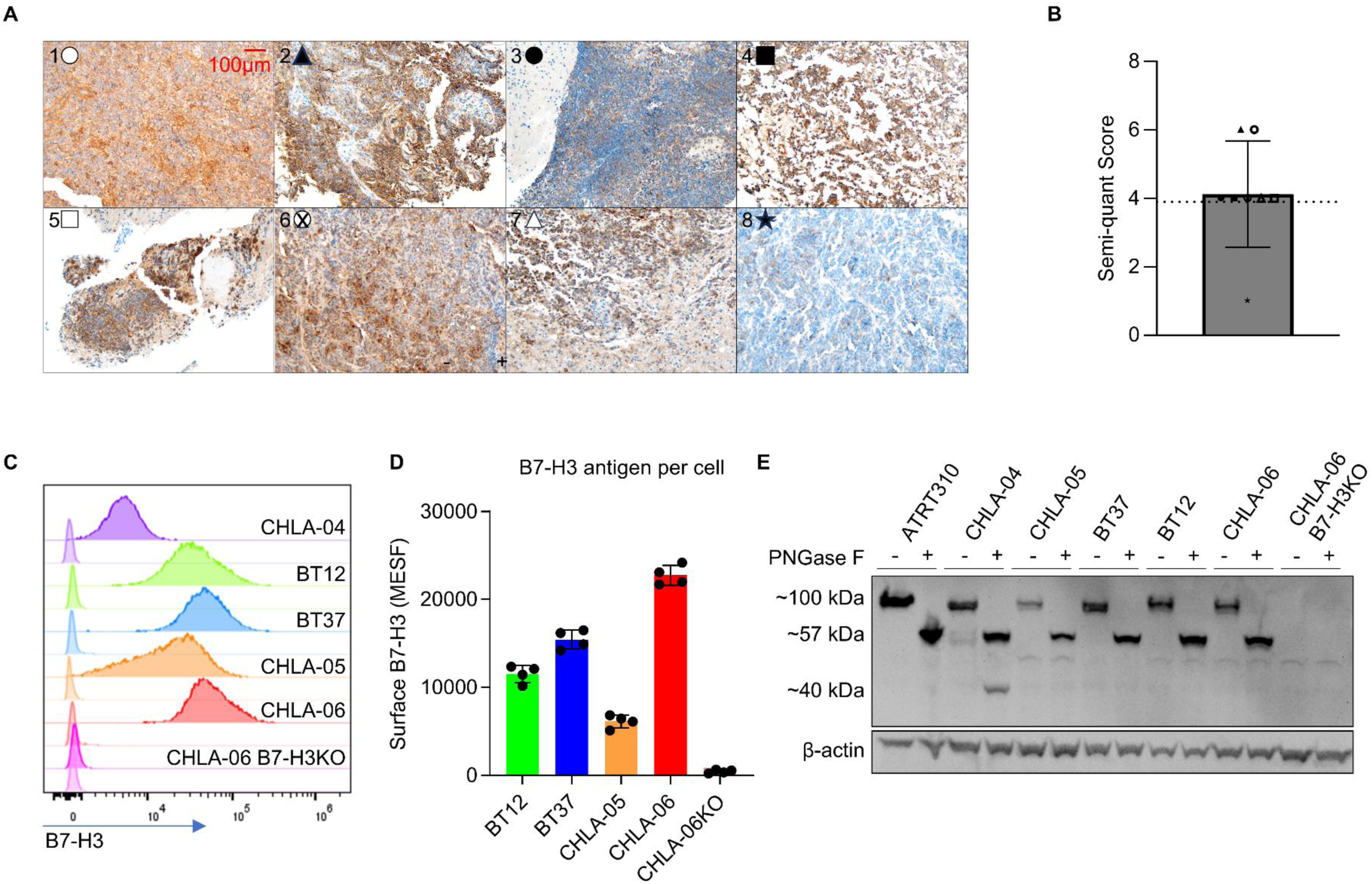
Patient-derived AT/RT cell lines and primary tumors express high amounts of B7-H3. (A) Immunohistochemistry (IHC) staining for B7-H3 on FFPE primary tumor tissue, and (B) semi-quantitative scoring of B7-H3 expression. 20X magnification, n=8. (C) AT/RT cell line B7-H3 expression. Isotype-stained samples are shown in the histograms directly below each B7-H3-stained sample. (D) B7-H3 surface molecules per cell quantified by molecules of equivalent soluble fluorochrome (MESF) flow cytometry assay. Each dot representative of technical replicates. (E) Total B7-H3 protein expression with or without deglycosylation assessed by western blot.

There are two known mRNA isoforms of human CD276 (2Ig and 4Ig) resultant from exon duplication^19,21^, but protein expression of the 2Ig product in tumor and normal tissue is minimal^12,22^. B7-H3 is highly glycosylated, which makes precise measurement of protein mass difficult^12,15,20^. When whole protein lysates from AT/RT cell lines were blotted for B7-H3, we observed a band at ∼100 kDa instead of the predicted 4Ig mass of 57 kDa (Figure S1B). To interrogate whether AT/RTs selectively express one isoform, whole protein lysates from AT/RT cell lines were digested with PNGase F to remove all N-linked glycosylation prior to B7-H3 detection. Most digested B7-H3 protein bands migrated at the predicted 4Ig mass (57 kDa), suggesting that AT/RT generally expresses exclusively the 4Ig isoform. Uniquely, undigested and digested CHLA-04 lysate reveals low-level 2Ig expression, with bands migrating at ∼60 kDa and the predicted mass of 34 kDa, respectively (Figure 1E). Our expression data corroborate previous reports of B7-H3 positivity in AT/RT cell lines and primary patient tissue^9,10^, with predominance of 4Ig isoform expression.

### Anti-B7-H3 CARs can be efficiently engineered into healthy donor-derived primary NK cells

To facilitate NK-cell target binding, we included a characterized humanized scFv extracellular domain with validated B7-H3 specificity in our previously described CAR backbone^34,35,41^. We generated mutants from the parent scFv sequence known to modify binding affinity (277.4-0.92nM)^34^ and cloned these into CAR constructs containing a 2B4 (CD244) hinge, transmembrane, and signaling domain, and a CD3ζ activation domain (Figure 2A, Supplementary Table 2). Primary healthy donor peripheral blood mononuclear cell (PBMC)-derived NK cells were activated and expanded *in vitro.* All scFv variant CARs were able to be transduced into and expressed on primary NK cells with high efficiency (Figure 2B,C). CAR transduced NK cells expanded comparably to untransduced (UTD) cells over 17 days of *in vitro* culture post-transduction (Figure 2D). Interestingly, activated and expanded unmodified NK cells expressed surface B7-H3 (Figure 2E,F), but anti-B7-H3 CAR-NK cells did not have detectable B7-H3 surface expression. When co-cultured with B7-H3-expressing CHLA-06 AT/RT cells, all anti-B7-H3 CAR-NK cells showed antigen-specific degranulation by CD107a staining, without differences related to scFv affinity (Figure 2G).

**Figure 2.**
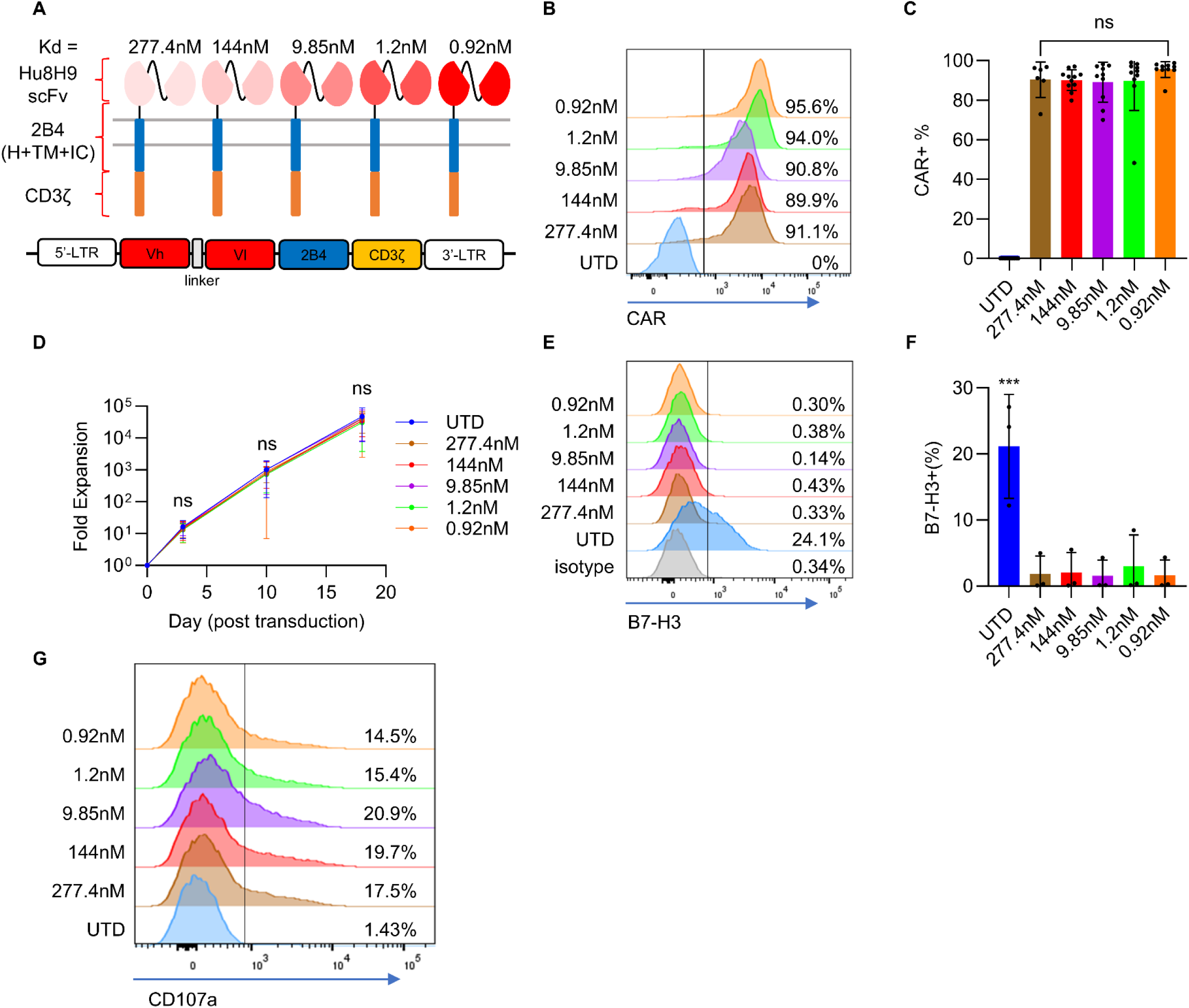
B7-H3 targeting CAR-NK cells can be efficiently expressed on primary human NK cells and are functional. (A) CARs consisting of Hu8H9 scFv and various affinity mutants, 2B4 (CD244) hinge (H), transmembrane (TM), and intracellular domains (IC), and CD3ζ signaling domain. (B) Representative primary NK-cell CAR expression. (C) CAR expression (n=4 healthy donors, 4-9 separate experiments). (D) Fold expansion of CAR-NK cells starting at day of transduction (n=4 healthy donors). (E) Representative CAR transduced NK-cell B7-H3 expression 2-4 days post-transduction. (F) B7-H3 expression on CAR-NK cells. (n = 3 healthy donors). *** = p<0.001 UTD vs. all others. (G) Representative measurement of CAR-NK cell degranulation post 2-hour co-culture with CHLA-06 AT/RT cells detected by CD107a staining.

### Anti-B7-H3 CAR-NK cells efficiently kill AT/RT *in vitro*

To assess the antigen-specific functionality of our CAR-NK cells, we co-cultured NK cells with ffLuc expressing target AT/RT cell lines at various E:T ratios. All CAR-NK cells had increased antigen-dependent cytotoxicity against B7-H3-expressing target cells as compared to UTD NK cells (Figure 3A). In this 18-hour co-culture assay, there was no correlation between scFv affinity and CAR-NK cell cytotoxicity to antigen-expressing tumor cells. UTD NK cells had roughly equivalent cytotoxicity against all target cell lines, with the exception of BT37 (TYR subtype) which was highly susceptible to NK-cell attack (Figure S2). In all co-culture assays, CAR expression significantly increased target cell killing if B7-H3 expression was present (Figure 3A). To assess longer-term cytotoxic potential, we co-cultured NK cells with AT/RT cell line targets expressing nuclear-localized GFP (NLS-eGFP) at a 1:1 E:T ratio over eight days. The culture was boosted with additional target cells every 48 hours. We again observed in this long-term assay that CAR-expressing NK cells killed AT/RT target cells in an antigen-specific manner, without affinity-dependent differences (Figure 3B,C). In summary, anti-B7-H3 CAR-NK cells have robust antigen-specific cytotoxicity against AT/RT, without detectable differences in cytotoxicity between CAR scFv affinities.

**Figure 3.**
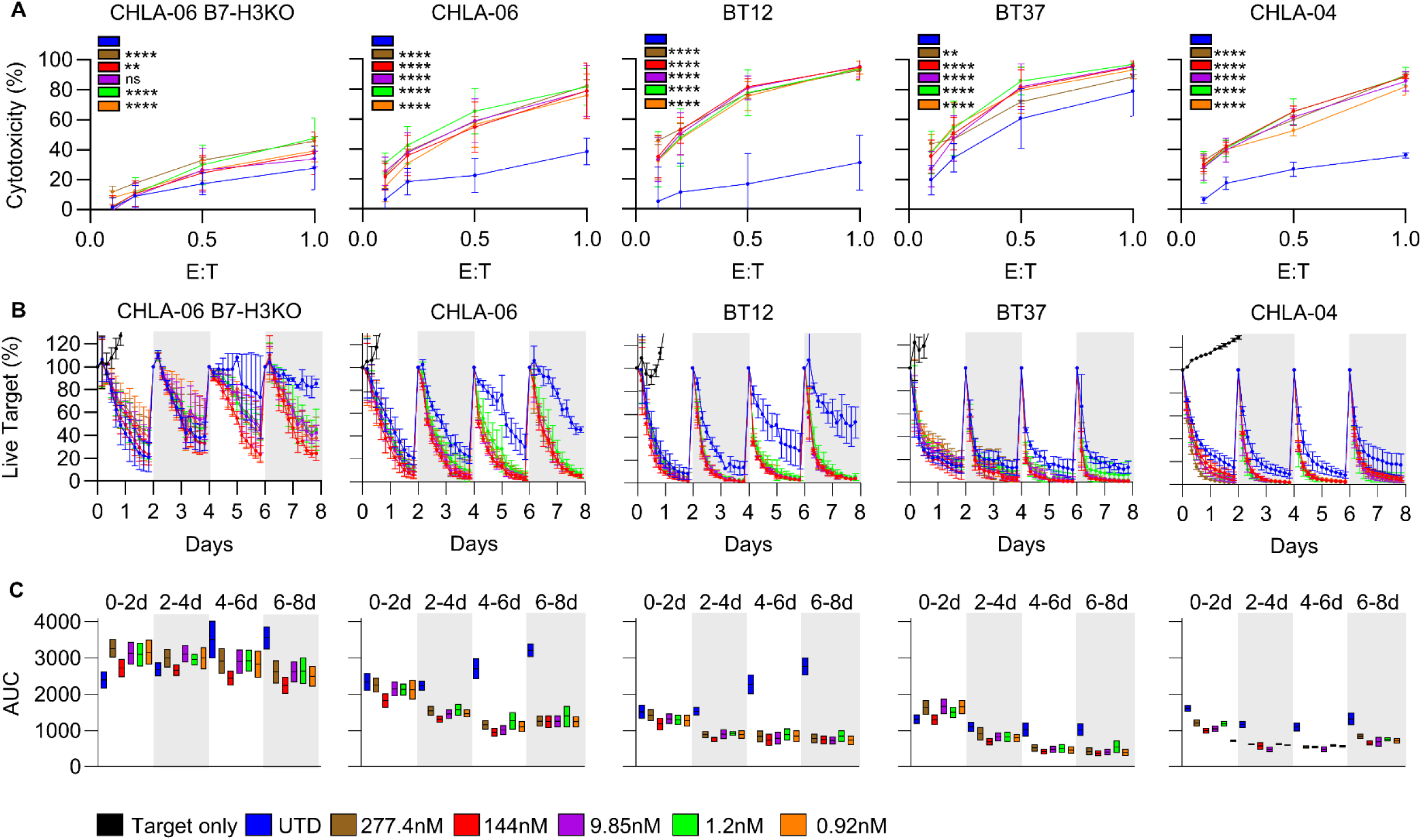
B7-H3-targeted CAR-NK cells show antigen-specific cytotoxicity against AT/RT cell lines *in vitro*. (A) NK-cell cytotoxicity versus indicated AT/RT cell lines after 18-hour co-culture relative to target only (n=4 donors, 2 donors for CHLA-04). *= significance of mean difference across all E:T ratios of each CAR compared to UTD. ** = p<0.01, **** = p<0.0001 (B) NK cells co-cultured with nuclear-localized eGFP (NLS-eGFP) expressing AT/RT target cell with target cell and IL-2 repletion every 48 hours. Live cells imaged every 4 hours. Percentage of live target cells was normalized to each stimulation with target cells. (C) Area Under Curve (AUC) analysis of data from (B) at each rechallenge timepoint, error bars showing 95% CI.

### Intratumoral delivery of anti-B7-H3 CAR-NK cells clears disease in an orthotopic xenograft model of human AT/RT

To test *in vivo* efficacy of anti-B7-H3 CAR-NK cells, we established orthotopic xenografts of CHLA-06.ffLuc (MYC subtype) through an implanted guide screw to facilitate repeat intratumoral delivery of therapeutic NK cells^42,43^. Animals were serially monitored for tumor growth and overall condition (Figure 4A). We opted to use a standard 144nM affinity CAR for *in vivo* treatment. UTD NK cells did not control tumor growth when compared to PBS-treated controls (Median survival = 16.5 and 20d, respectively), whereas anti-B7-H3 CAR-NK cells were able to control tumor growth in a majority of treated mice (Median survival = undefined). Responding animals remained tumor-free until experimental end (Figure 4B,C,D). Upon histological analysis of brains at humane experimental endpoints, we confirmed that treatment-resistant tumors did not lose target B7-H3 expression in any condition (Figure 4E, Figure S3A). Both UTD and CAR-NK cells were detectable in tumors up to six days post-treatment (Figure 4E). In the mice that were cured, we noted that the effect of CAR-NK cell treatment was evident after the initial NK-cell injection (Figure 4A,B). To interrogate the anti-tumor kinetics surrounding the NK-cell infusion, we performed a separate parallel tumor treatment experiment with identical conditions and planned necropsy at 48 hours post NK cell treatment (Figure S3B). At this timepoint, we observed CAR-NK treated mice had increased amounts of reactive gliosis in the primary tumor site shown by increased F4/80 staining, whereas UTD NK treated mice had larger, viable areas of tumor (Figure S3B). We observed the location of NK cells (whether UTD or CAR-NK) to be perivascular, periventricular, and peritumoral with minimal infiltration into the normal brain parenchyma (Figure 4E, Figure S3B). Ventricular tumor seeding was present in all mice (Figure S3C).

**Figure 4.**
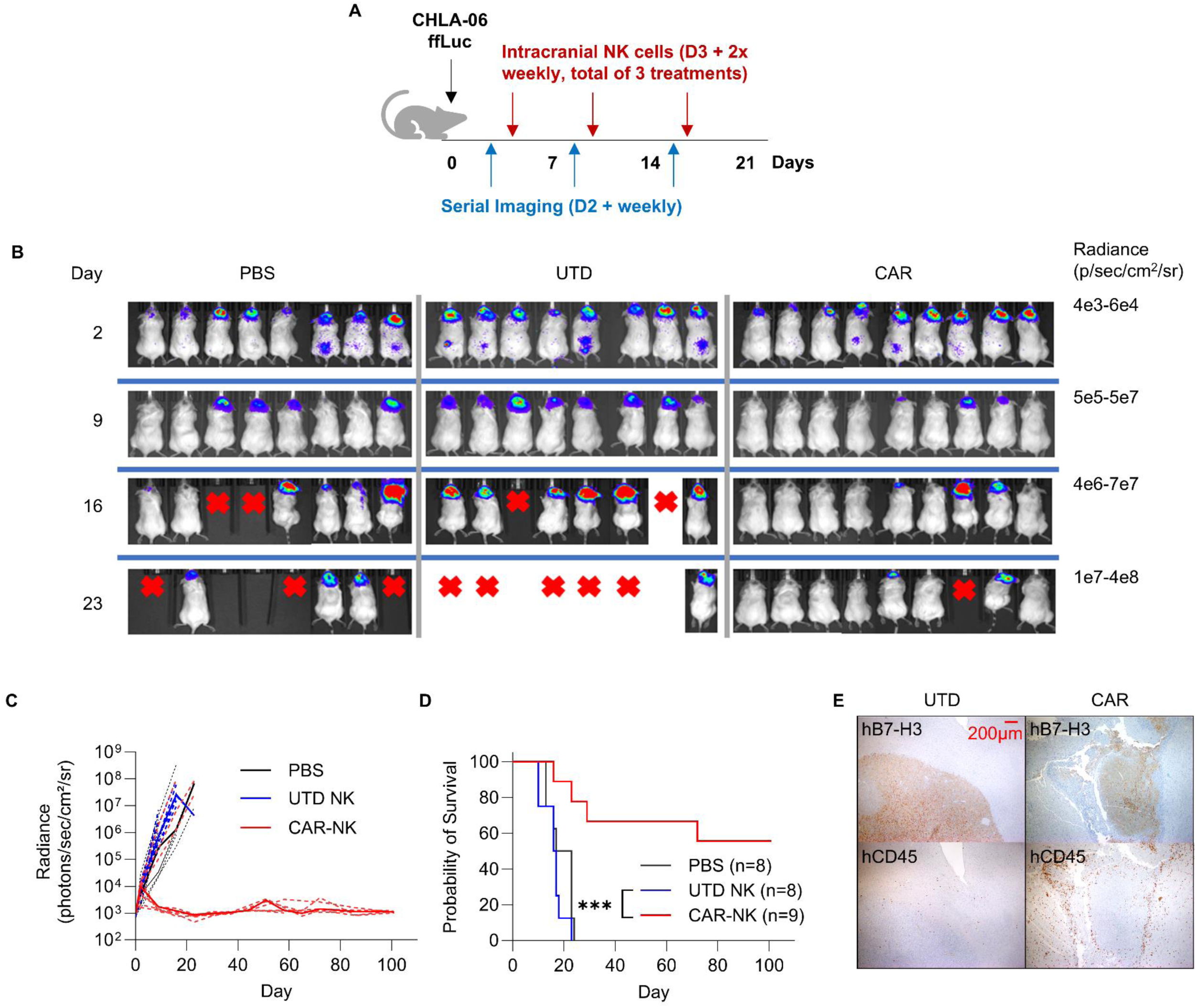
Anti-B7-H3 CAR-NK cells significantly improve survival and reduce tumor burden when delivered to orthotopic xenografts of CHLA-06 AT/RT. (A) CHLA-06.ffLuc (25k cells) were injected into caudate putamen of NSG mice at day 0. Randomization/cohorting was performed at day 2, and intratumoral NK cell treatment (3 million cells per treatment) was started on day 3 and repeated on days 10 and 17. (B) Tumor radiance measured by IVIS Spectrum bioluminescent imaging and analyzed using Living Image software. (C) tumor radiance over time of CHLA-06.ffLuc bearing mice (n=8-9 per condition). (D) Kaplan-Meier survival curve. ***= p<0.001. (E) Brain tissue harvested at endpoint (D23 post-tumor, D6 post-third NK-cell treatment for both samples), representative slides from UTD NK treated and CAR-NK treated mice stained for human B7-H3 and CD45. 40X magnification.

### Intracranioventricular (ICV) delivery of anti-B7-H3 CAR-NK cells has anti-tumor activity in a disseminated AT/RT tumor model

To test anti-B7-H3 CAR-NK cell control of AT/RT with leptomeningeal spread, we developed a BT12 disseminated tumor model. We established tumor intracerebroventricularly (ICV), verified tumor take by bioluminescent imaging, then administered PBS, UTD, or CAR-NK cells by ICV injection (Figure 5A, Figure S4). While PBS control and UTD NK treated animals experienced accelerated tumor spread and early death due to widespread disease (Median survival for both = 27d), ICV CAR-NK treatment slowed tumor growth (Median survival = 72d) and prolonged survival (Figure 5B). We also observed delayed weight loss in the CAR-NK treated mice (Figure 5C). Upon examination of brain and spinal cord tissue at endpoint, we observed prominent intracranial disease in PBS and UTD NK treated mice, with lower overall intracranial disease burden in CAR-NK treated animals (Figure 5D,E). Strikingly, some CAR-NK treated animals had no detectable brain tumors. Regardless of treatment condition, mice with intracranial disease had tumors present in ventricles and in the leptomeningeal space surrounding the brain. All mice had disseminated spinal cord tumors (Figure 5A,D,E, Figure S5). Staining of tissue for hCD45 revealed increased NK cell numbers in the CAR-NK treatment group (Figure 5F). CAR-NK cells were enriched in both intracranial and spinal tumors when compared to UTD NK cells. In summary, anti-B7-H3 CAR-NK cell treatment effectively prolonged survival in an AT/RT model of leptomeningeal metastasis, with CAR-NK cell infiltration and enrichment at sites of disease.

**Figure 5.**
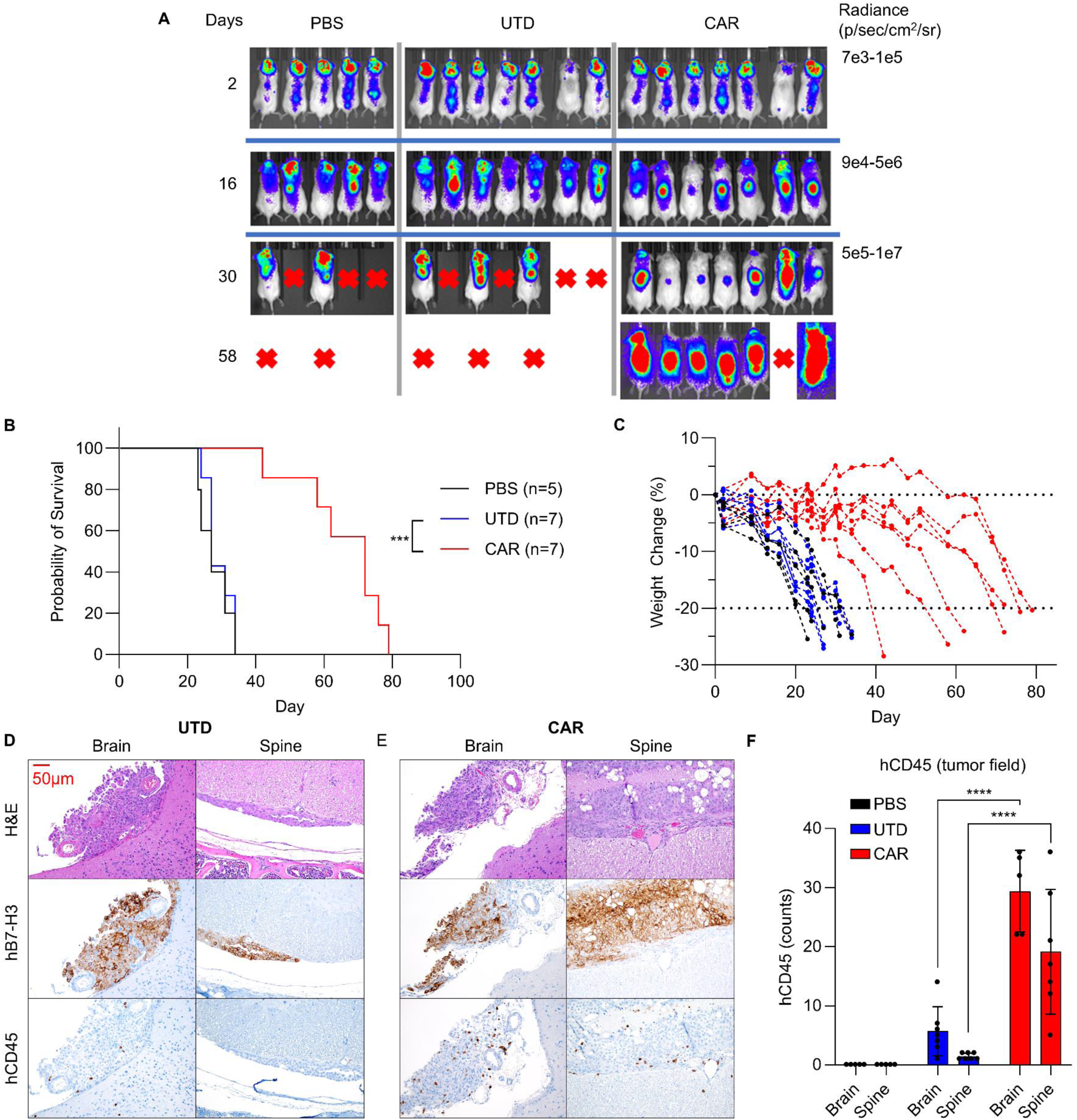
Intracerebroventricular (ICV) CAR-NK cell treatment of craniospinal disseminated BT12 AT/RT significantly improves survival. (A) BT12.ffLuc (100k cells) were injected into the right lateral ventricle of NSG mice at day 0. Randomization/cohorting was performed at day 2, and ICV NK cell treatment was started on day 3 and repeated on days 10 and 17. (B) Kaplan-Meier survival curve, and (C) percent weight change over time of ICV BT12.ffLuc bearing mice. Each dotted line represents an individual animal. Representative histology of endpoint brain and spinal tissue from (D) UTD NK treated, and (E) CAR-NK treated mice with corresponding human B7-H3 and CD45 IHC stains. Animal endpoints = 31, 62 days post-tumor injection, respectively. 200X magnification. (F) Quantification of tumor associated human NK cells. n = 5-7 per condition.

## Discussion

AT/RT remains a difficult malignancy to cure, with patients suffering from treatment resistance, relapse, and treatment-related morbidities. We show in primary samples and established AT/RT cell lines that the tumors express high levels of the pan-cancer antigen B7-H3 (CD276). This observation validates prior studies that previously have shown B7-H3 expression to be tumor-specific and safe to target with engineered T-cells^10^. We efficiently engineered healthy donor peripheral blood-derived NK cells with multiple functional B7-H3 targeting CARs, all of which stimulate antigen-dependent cytotoxicity against AT/RT *in vitro*. Other preclinical studies testing B7-H3 targeting CAR-NK cells^44,45^ as glioma therapy and clinical trials testing B7-H3 targeting CAR-T cells^11,46^ in pediatric CNS disease use an scFv derived from the MGA271 (enoblituzumab) monoclonal antibody^47^. We instead chose to incorporate a B7-H3 binding scFv derived from the humanized 8H9 (omburtamab) monoclonal antibody^34,41^ for development and functional testing of CAR-NK cells due to its well characterized binding epitope and supporting affinity data. We aligned our NK activation and expansion protocol to closely resemble that of cell products in an ongoing phase 1 clinical trial evaluating *ex vivo* expanded, TGF-β1 imprinted, universal donor NK cells in recurrent pediatric brain tumors, including AT/RT (NCT05887882)^48^. Imprinting of activated NK cells with low concentrations of TGF-β1 has previously been shown to generally enhance cytotoxicity against solid tumors^49^. Thus, we maintained TGF-β1 supplementation during CAR-NK cell manufacturing as standard.

We have previously shown CAR scFv affinity dependent effects in the context of targeting CD123 in acute myeloid leukemia^50^, hence we explored the possibility of similar effects in conditions tested in this study. Evaluation of CAR expression, CAR-NK cell antigen-dependent activation, and CAR-NK cell directed cytotoxicity illuminated no differences related to CAR binding affinity. We therefore conclude that, in the context of targeting B7-H3-expressing AT/RT, CAR scFv affinity has no impact on CAR-NK cell functionality. Given the binding complexity of NK cell interactions with target cells, cellular avidity is likely a more reliable predictor of anti-tumor activity than solely considering CAR affinity^51^.

We confirm that for the majority of AT/RT, B7-H3 is expressed as its 4Ig isoform. Knowledge of the 8H9 binding epitope^34^ and the 4Ig structure^21^, allows presumption that our CARs would face duplicate binding opportunities in each of the IgV regions of 4Ig B7-H3. This interaction nuance, unique to B7-H3 structural duplication, is very likely consequential and should underpin further study of cellular avidity, binding kinetics, and the determinant factors of productive immunological synapse formation when B7-H3 is used as a cancer-specific target antigen.

Our CAR-NK cells persist in BT12 xenografts for up to 62 days after injection as evidenced by human CD45 IHC performed on endpoint tissue. In prior work, we observed a lack of NK cell *in vivo* persistence unless the cells were engineered with constitutive cytokine support^35^. Yet, the CAR-NK cells in this study persisted in tumor for weeks after CSF administration. It is unknown how the AT/RT tumor environment promotes NK-cell persistence, and the durability of our B7-H3 CAR-NK cells *in vivo* may undergird the remarkable improved survival we observe in two aggressive intracranial AT/RT models. While we were able to cure CHLA-06 bearing mice with intratumoral injections of CAR-NK cells, complete disease clearance was not observed in our BT12 ICV model. The disseminated nature of the tumor and NK cells in the latter model may contribute to the observed incomplete anti-tumor response, with NK cell suppression by tumor inhibitory ligands also likely. A better understanding of how the tumor microenvironment affects NK-cell killing of AT/RT is a major focus of our future investigations.

In this study, we demonstrate *in vivo* utility of healthy donor peripheral blood derived, cryopreserved, B7-H3 targeting CAR-NK cells as a monotherapy in multiple preclinical models of AT/RT without exogenous cytokine support, an intentional mimic of how these cells can be used clinically. We validate massive expansion and cryopreservation of healthy donor NK cells^48^, allowing for ready infusion and reinfusion to patients.

Installation of Ommaya reservoirs to allow for direct CNS access has become common for patients enrolled in therapeutic clinical trials, allowing for repeated injections of chemotherapy or cell therapy agents in outpatient settings^52^. Healthy donor-derived, *ex vivo* activated primary NK cells are clinically safe and are currently under investigation with direct CNS delivery to pediatric brain tumor patients (NCT05887882)^48^. We demonstrated efficacy of both intratumoral and ICV delivered NK cells in distinct models of AT/RT. We thus believe that the addition of a tumor-specific CAR to transferred NK cells will be beneficial and enhance therapeutic response in this vulnerable population.

## Ethics

Animal experiments were done in accordance with protocol approved by Johns Hopkins Animal Care and Use Committee (IACUC).

## Authorship Statement

Conceptualization: J.C., E.H.R., and C.L.B. Performance of experiments: J.C., S.C., N.J.H., M.T.Z., D.G.J., S.G., R.R., S.C.V., and A.A. Analysis of results: J.C., C.G.L, E.H.R., and C.L.B.

Drafting of manuscript: J.C. E.H.R., and C.L.B. Manuscript review: All authors.

## Data availability

All data associated with this study are present in the paper and Supplementary Materials. Any additional data or raw files are available upon request.

## Acknowledgements

We would like to thank Dr. Charles Eberhart, Dr. Robyn Gartrell, and Dr. Dean Lee for helpful discussion. We would also like to thank Dr. Elizabeth Jaffee and Dr. Giorgio Raimondi for being a part of J.C.’s thesis committee and contributing helpful discussion.

