## Supplemental Material for "B7-H3-targeted natural killer cells effectively kill atypical teratoid / rhabdoid tumors and extend survival in orthotopic xenografts"

**Supplementary Figure 1**

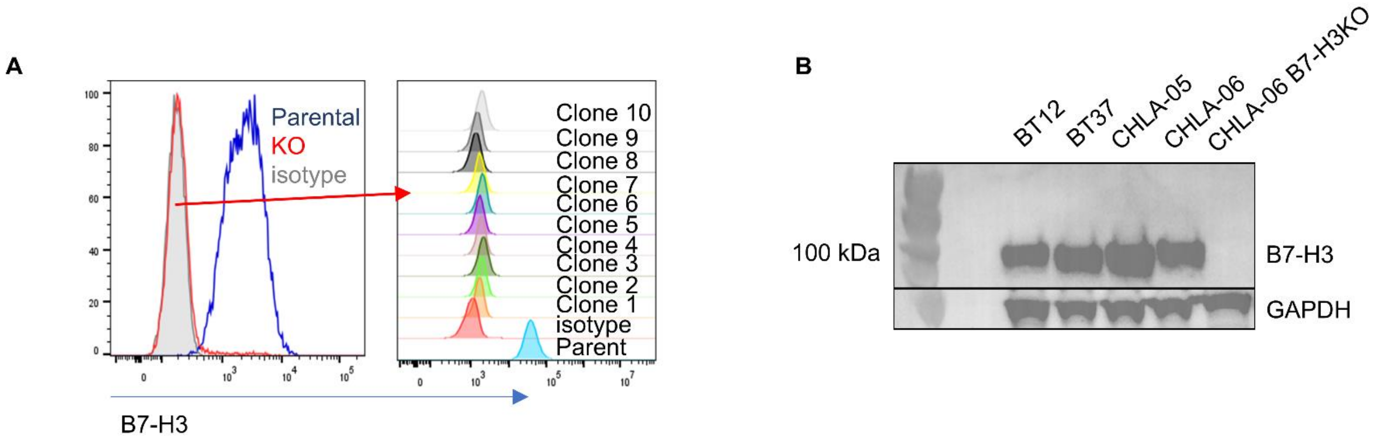


**Figure S1. CD276 CRISPR-Cas9 KO of CHLA-06 effectively generated viable clones.** (A) KO efficiency of *CD276* in CHLA-06 post RNP electroporation assessed by flow cytometry. Single cell clones were isolated and pooled. (B) Western blot of total B7-H3 from representative AT/RT cell lines shows that endogenous B7-H3 migrates at ~100 kDa instead of the predicted 57 kDa for the 4Ig protein form and that there is no evident protein expression in the pooled KO clones.

**Supplementary Figure 2**

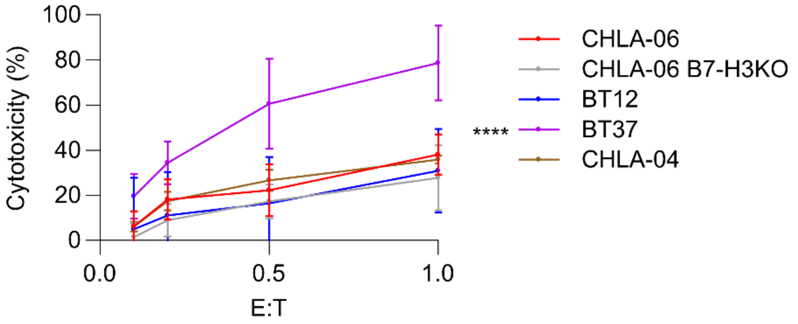


**Figure S2. Innate cytotoxicity of UTD NK cells against various AT/RT targets.** Cytotoxicity data from Figure 3A revisualized to compare innate cytotoxicity of UTD NK cells against all targets. *= significance between BT37 vs. all other target conditions (n=4 healthy donors, 2 donors for CHLA-04). **** = p<0.0001.

**Supplementary Figure 3**

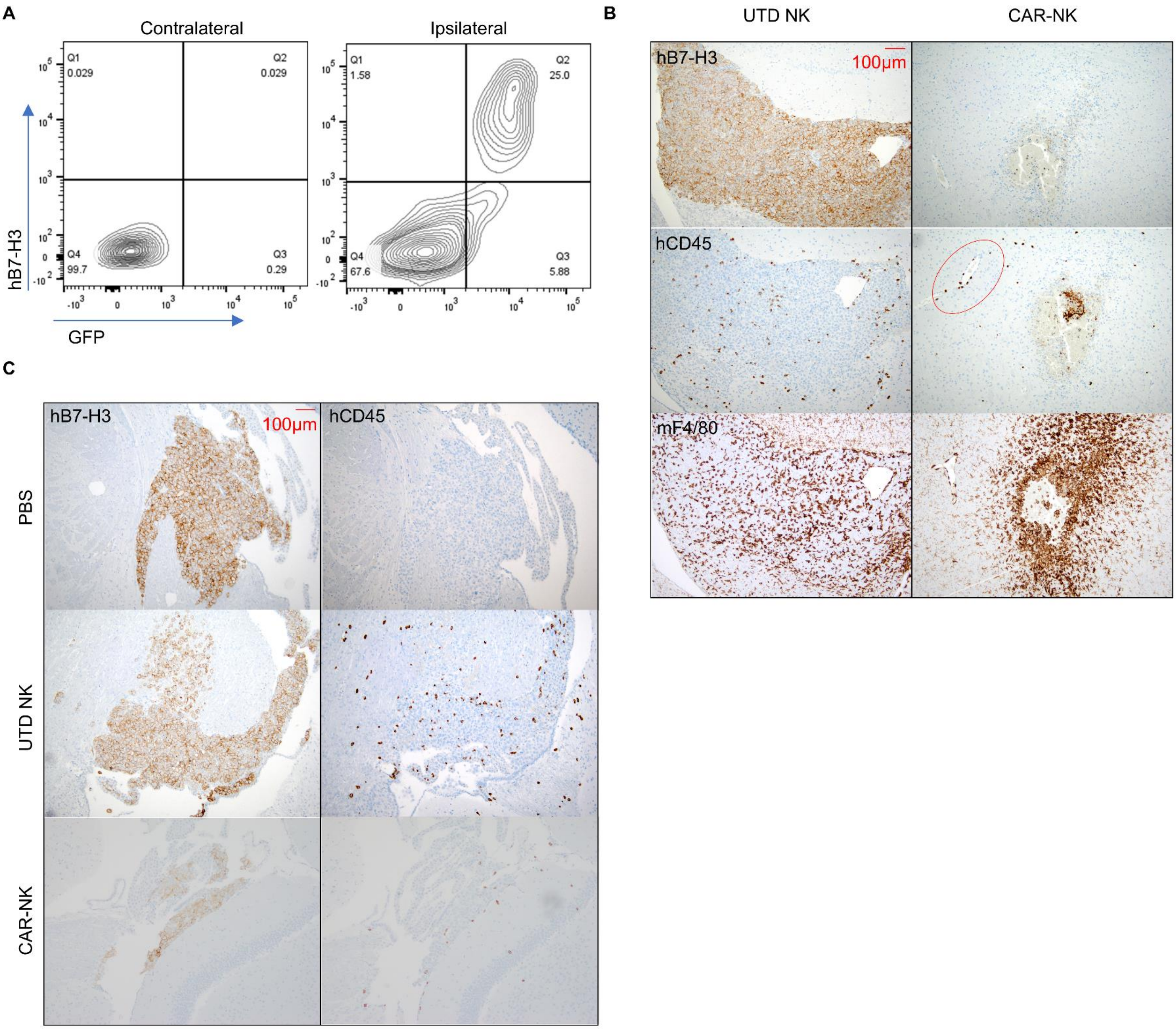


**Figure S3. B7-H3 expression is retained in orthotopic xenografts and transferred human NK cells can be detected by IHC.** (A) Brain tissue from a PBS-treated CHLA-06.ffLuc.GFP bearing NSG mouse taken at endpoint (D23 post-tumor). Contralateral and ipsilateral brain was bisected from whole brain and dissociated into single cells for flow cytometry staining. (B) Brain tissue harvested 48-hours post-NK-cell treatment (D5 post tumor implantation) from UTD NK treated and CAR-NK treated mice stained for human B7-H3, human CD45, and mouse F4/80. 100X magnification. Perivascular NK cells circled in red. (C) Ventricular seeding of CHLA-06 tumor observed at 48-hours post-NK-cell treatment, 5 days post tumor injection. 100X magnification.

**Supplementary Figure S4**

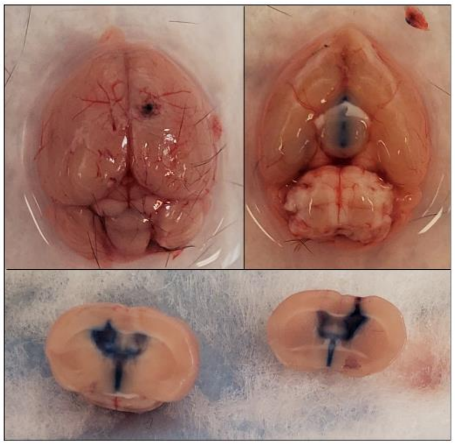

**Figure S4. ICV injection facilitates ventricular dissemination of injected material.** Representative ICV injection (right lateral ventricle) of Evans Blue dye into naïve mouse 1-hour post-injection.

**Supplementary Figure S5**

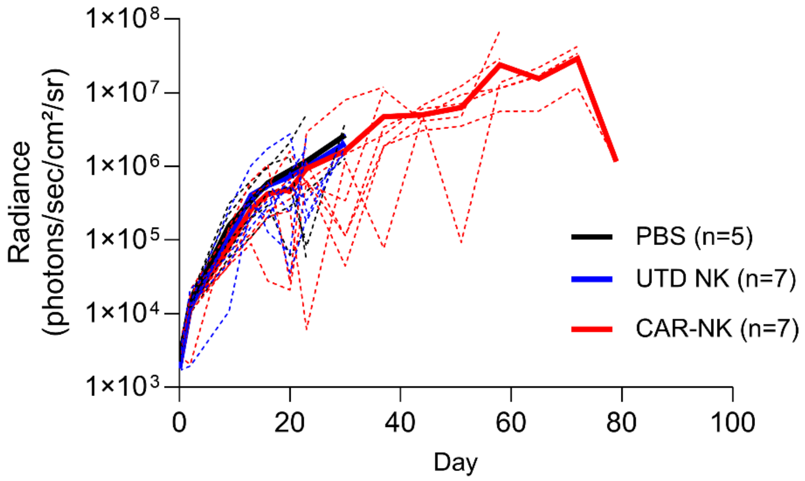


**Figure S5. CAR-NK treatment did not significantly reduce total tumor burden in disseminated BT12.** Tumor radiance over time of ICV BT12 bearing mice, quantified across whole bodies. Dotted lines show individual values and solid lines show median (n=5-7 per condition).

**Supplementary Table 1**

| Primer/guide | Application | Sequence (5'-3') |
|---|---|---|
| Vector.2B4.FOR | In-Fusion | CAGGATTGCCAGAATGCCCAC |
| Vector.SP.REV | In-Fusion | AGAATGGGCGCCTGTGG |
| Hu8H9.FOR | In-Fusion | ACAGGCGCCCATTCTCAGGTTCAGCTGGTTCAGTCT |
| Hu8H9.REV | In-Fusion | ATTCTGGCAATCCTGGGATCCGCCGCCACC |
| a262c_g263c | SDM | TGGAACTGAGCAGCCTGAGACCCGAGGATACCG |
| a565c_g566c_as | SDM | GGCGGGGATGCCGGGGATAGACTGGCTGG |
| a565c_g566c | SDM | CCAGCCAGTCTATCCCCGGCATCCCCGCC |
| a262c_g263c_as | SDM | CGGTATCCTCGGGTCTCAGGCTGCTCAGTTCCA |
| g305c_as | SDM | TAGGCAAACCAAGTTGCGGTGGTCTGTCTGG |
| g305c_ | SDM | CCAGACAGACCACCGCAACTTGGTTTGCCTA |
| a167g | SDM | TTCCCTGGCGACGGCAGCACCCAGTAC |
| a167g_as | SDM | GTACTGGGTGCTGCCGTCGCCAGGGAA |
| R87G.f | SDM | GAACTGAGCAGCCTGGGAAGCGAGGATACCG |
| R87G.r | SDM | CGGTATCCTCGCTTCCCAGGCTGCTCAGTTC |
| A97T.f | SDM | CCGTGTACTTCTGCACCAGACAGACCACC |
| A97T.r | SDM | GGTGGTCTGTCTGGTGCAGAAGTACACGG |
| G129S.f | SDM | GGTGGCGGAGGAAGTAGCGGAGGC |
| G129S.r | SDM | GCCTCCGCTACTTCCTCCGCCACC |
| S163G.f | SDM | CCAGCCAGAGCATCGGCGACTACCTGTAC |
| S163G.r | SDM | GTACAGGTAGTCGCCGATGCTCTGGCTGG |
| pSFG-F | Sequencing | AGCTTGGATACACGCCGCC |
| SFG-R2 | Sequencing | GCTCGTACTCTATAGGCTTCA |
| 276.Sgo.f2 | KO PCR | CAGTTGCAGCCTTCCCCA |
| 276.Sgo.r1 | KO PCR | CACAGTGTATTCAAGAAATAGCACGA |
| CD276 sgRNA1 | CD276 KO | CTCACAGGAAGATGCTGCGT |
| CD276 sgRNA2 | CD276 KO | GCACTGTGGTTCTGCCTCAC |
| CD276 sgRNA3 | CD276 KO | GGGACAGTGATTGTGGCAGT |

**Supplementary Table 2**

| Sequence | Source | aa Sequence |
|---|---|---|
| 277.4 nM (scFv) | Ref 34 and this study | MDWIWRILFLVGAATGAHSQVQLVQSGAEVVKPGASVKLSCKTSGYTFTNYDINWVRQRPGQGLEWIGWIFPGDGSTQYNEKFKGKATLTTDTSTSTAYMELSSLGSEDTAVYFCTRQTTATWFAYWGQGTLVTVSSGGGGSGGGGSSGGGSEIVMTQSPATLSVSPGERVTLSCRASQSIGDYLYWYQQKSHESPRLLIKYASQSISGIPARFSGSGSGSEFTLTINSVEPEDVGVYYCQNGHSFPLTFGQGTKLELKRSGGGGS |
| 144 nM (scFv) | Ref 34 and this study | MDWIWRILFLVGAATGAHSQVQLVQSGAEVVKPGASVKLSCKASGYTFTNYDINWVRQRPEQGLEWIGWIFPGDGSTQYNEKFKGKATLTTDTSTSTAYMELSSLRSEDTAVYFCARQTTATWFAYWGQGTLVTVSSGGGGSGGGGSGGGGSEIVMTQSPATLSVSPGERVSLSCRASQSISDYLHWYQQKSHESPRLLIKYASQSISGIPARFSGSGSGSEFTLTINSVEPEDVGVYYCQNGHSFPLTFGQGTKLELKRSGGGGS |
| 9.85 nM (scFv) | Ref 34 and this study | MDWIWRILFLVGAATGAHSQVQLVQSGAEVVKPGASVKLSCKTSGYTFTNYDINWVRQRPGQGLEWIGWIFPGDGSTQYNEKFKGKATLTTDTSTSTAYMELSSLRSEDTAVYFCARQTTATWFAYWGQGTLVTVSSGGGGSGGGGSGGGGSEIVMTQSPATLSVSPGERVTLSCRASQSISDYLYWYQQKSHESPRLLIKYASQSISGIPARFSGSGSGSEFTLTINSVEPEDVGVYYCQNGHSFPLTFGQGTKLELKRSGGGGS |
| 1.2 nM (scFv) | Ref 34 and this study | MDWIWRILFLVGAATGAHSQVQLVQSGAEVVKPGASVKLSCKTSGYTFTNYDINWVRQRPGQGLEWIGWIFPGDDSTQYNEKFKGKATLTTDTSTSTAYMELSSLRPEDTAVYFCARQTTGTWFAYWGQGTLVTVSSGGGGSGGGGSGGGGSEIVMTQSPATLSVSPGERVTLSCRASQSISDYLYWYQQKSHESPRLLIKYASQSIPGIPARFSGSGSGSEFTLTINSVEPEDVGVYYCQNGHSFPLTFGQGTKLELKRSGGGGS |
| 0.92 nM (scFv) | Ref 34 and this study | MDWIWRILFLVGAATGAHSQVQLVQSGAEVVKPGASVKLSCKTSGYTFTNYDINWVRQRPGQGLEWIGWIFPGDDSTQYNEKFKGKATLTTDTSTSTAYMELSSLRSEDTAVYFCARQTTGTWFAYWGQGTLVTVSSGGGGSGGGGSGGGGSEIVMTQSPATLSVSPGERVTLSCRASQSISDYLYWYQQKSHESPRLLIKYASQSISGIPARFSGSGSGSEFTLTINSVEPEDVGVYYCQNGHSFPLTFGQGTKLELKRSGGGGS |
| 2B4 (hinge, TM, cytoplasmic) + CD3ζ | Ref 35 and this study | QDCQNAHQEFRFWPFLVIIVILSALFLGTLACFCVWRRKRKEKQSETSPKEFLTIYEDVKDLKTRRNHEQEQTFPGGGSTIYSMIQSQSSAPTSQEPAYTLYSLIQPSRKSGSRKRNHSPSFNSTIYEVIGKSQPKAQNPARLSRKELENFDVYSGAGRVKFSRSADAPAYQQGQNQLYNELNLGRREEYDVLDKRRGRDPEMGGKPQRRKNPQEGLYNELQKDKMAEAYSEIGMKGERRRGKGHDGLYQGLSTATKDTYDALHMQALPPR* |

**Supplementary Table 3**

| Antibody/stain | Clone | Manufacturer | Cat# | Application | Dilution |
|---|---|---|---|---|---|
| anti-CD276 APC | MIH42 | Biolegend | 351006 | FC | 1:50 |
| anti-CD276 | D9M2L | Cell Signaling | 14058 | IHC | 1:100 |
| anti-CD276 | SP206 | Abcam | ab227670 | WB | 1:1000 |
| anti-β actin | C4 | Santa Cruz | sc-47778 | WB | 1:10000 |
| anti-GAPDH | 6C5 | Invitrogen | AM4300 | WB | 1:5000 |
| anti-His APC | J095G46 | Biolegend | 362605 | FC | 1:20 |
| anti-His PE | J095G46 | Biolegend | 362603 | FC | 1:20 |
| anti-CD107a APC | H4A3 | Biolegend | 328620 | FC | 1:800 |
| anti-CD45 PE | HI30 | Biolegend | 304058 | FC | 1:20 |
| anti-CD3ε PE | 145-2C11 | BD Biosciences | 553063 | FC | 1:20 |
| anti-CD56 BV421 | NCAM16.2 | BD Biosciences | 562751 | FC | 1:20 |
| anti-CD247 | 1D4 | BD Biosciences | 556366 | WB | 1:1000 |
| anti-CD247 (pY142) | K25-407.69 | BD Biosciences | 558402 | WB | 1:1000 |
| Live/Dead Fixable Violet | | Invitrogen | L34963 | FC | 1:1000 |
| Live/Dead Fixable Green | | Invitrogen | L34969 | FC | 1:1000 |

**Supplementary Table 4**

| Number | Age | Sex | Surgery | Location | B7-H3 Intensity | B7-H3 Extent | Score |
|---|---|---|---|---|---|---|---|
| 1 | 5yrs | F | Primary | Left frontal | 2 | 3 | 6 |
| 2 | 3yrs | F | Primary | Cerebellum | 2 | 3 | 6 |
| 3 | 6mos | F | Primary | Right lateral ventricle | 2 | 2 | 4 |
| 4 | 6mos | F | Primary | Cerebellum | 2 | 2 | 4 |
| 5 | 1yrs | M | Primary | Left temporal | 2 | 2 | 4 |
| 6 | 4yrs | F | Primary | Right frontotemporal | 2 | 2 | 4 |
| 7 | 3yrs | F | Primary | Right frontotemporal | 2 | 2 | 4 |
| 8 | 7mos | F | Primary | Cerebellum | 1 | 1 | 1 |

**Supplementary methods**

*Cell line culture and reporter cell line generation*

AT/RT cell lines BT12 (RRID:CVCL_M155), BT37 (RRID:CVCL_JL57), CHLA-04 (RRID:CVCL_0F38), CHLA-05 (RRID:CVCL_AQ41), and CHLA-06 (RRID:CVCL_AQ42) have previously been described[1,2]. BT12 was obtained through the Children's Oncology Group cell repository. BT37 was derived from a human xenograft originating at St. Jude Children's Research Hospital[3]. ATRT-310 was obtained from Seattle Children's Hospital Brain Tumor Resource Lab. CHLA-04, CHLA-05, and CHLA-06 were a kind gift from Dr. Anat Erdreich-Epstein. BT12 and BT37 were maintained in Roswell Park Medical Institute (RPMI, Gibco) medium supplemented with 10% fetal bovine serum (FBS, HyClone). ATRT-310, CHLA-04, CHLA-05, and CHLA-06 were maintained in neurobasal medium consisting of 7:3 DMEM:F12 (Gibco) containing 2% B-27 supplement without vitamin A (ThermoFisher, 12587010), 1% L-glutamine (ThermoFisher A2916801), 20 ng/mL epidermal growth factor (Peprotech AF-100-15), 20 ng/mL basic fibroblast growth factor (Peprotech 100-18B), and 5 μg/mL heparin (Sigma-Aldrich H3149). HEK293T (ATCC Cat# CRL-3216) cells used for viral production were maintained in Dulbecco's Modified Eagle Medium (DMEM, Gibco) complete with 10% FBS and were originally obtained from ATCC. 293Vec-RD114 and 293Vec-BaEV producer cells were maintained in DMEM with 10% FBS and were obtained from BioVec Pharma[4]. Cell lines were transduced with replication incompetent VSV-G pseudotyped retroviral vectors containing ffLuc-eGFP or NLS-eGFP reporters encoded in pSFG plasmids to enable target cell tracking as indicated. All transduced cell lines were sorted using a BD FACSMelody Cell Sorter and confirmed >90% positive for reporters by FACS and by using substrates for enzymes when applicable. Cell lines were authenticated (Johns Hopkins Genetic Resources Core Facility) and tested for mycoplasma (MycoAlert Detection kit, Lonza) when new cell lines were established after transduction and FACS sorting, and before cryopreservation.
